# Microgravity affects the nervous system and aging in *C. elegans* through reduced tactile stimulation

**DOI:** 10.64898/2026.02.16.706116

**Authors:** Atsushi Higashitani, Je-Hyun Moon, Jong-In Hwang, Nahoko Higashitani, Toko Hashizume, Ahmad Aisha Abu, Kazuki Ooizumi, Ibuki Sazuka, Yoshimitsu Hashizume, Masumi Umehara, Alfredo V. Alcantara, Ban-seok Kim, Timothy Etheridge, Nathaniel J. Szewczyk, Takaaki Abe, Jin I. Lee, Akira Higashibata

## Abstract

Space travel is becoming accessible, yet our understanding of how space environment and microgravity (µG) affect biology, physiology, and health remains incomplete. We investigated µG effects on neuromuscular development and aging in *Caenorhabditis elegans*. Nematodes in µG showed downregulation of genes related to synaptic signaling, dopamine response, locomotion, and cuticle development, with impaired synaptic vesicle dynamics, reduced motility, and shorter body lengths. Aged worms in µG showed decreased collagen gene expression, increased motor neuron defects, synaptic vesicle accumulation and decreased release, and mitochondrial morphology collapse in body wall muscles, indicating accelerated aging. MEC-4 mechanoreceptor was identified as a key mediator of µG-induced body length reduction and changes in extracellular matrix gene expression. µG conditions suppressed mechanoreceptor genes, suggesting multiple mechanosensory systems are affected. Physical stimulation through culture medium with small beads in space mitigated many µG-induced expression changes, including mechanoreceptors, neuromuscular defects, and aging-related phenotypes. These results highlight mechanical stimuli’s role in maintaining neuromuscular integrity during spaceflight and suggest restoring tactile input could counter health risks from reduced stimulation in long-term space missions.

**SIGNIFICACE:** We found that microgravity (µG) conditions suppress the expression of multiple mechanoreceptor genes in *Caenorhabditis elegans*, indicating that several mechanosensory systems are affected during spaceflight. Importantly, reintroducing physical stimulation by adding small beads to the culture medium in space partially reversed many of these µG-induced gene expression changes. This intervention also mitigated neuromuscular defects and aging-related phenotypes observed under µG conditions. Collectively, these findings underscore the essential role of mechanical stimuli in preserving neuromuscular integrity during space missions and suggest that restoring tactile input may be a promising strategy to counteract the health risks associated with reduced tactile stimulation during prolonged spaceflights.

## Introduction

Technological advances have enabled extended human habitation in outer spaces. However, microgravity (µG) can induce aging-like symptoms, such as osteoporosis and muscle atrophy 1,2. Multi-omics analysis of NASA GeneLab data revealed that mitochondrial stress was a central factor, and that biomarkers of aging and frailty were increased following spaceflight 3,4. In addition, spaceflight mice showed decreased expression of dopamine-related genes such as *TH, COMT, MaoA, and D1R*, a phenomenon not observed in tail suspension models 5,6. Therefore, understanding the neuromuscular decline induced by spaceflight not only supports space activities but may also enhance the quality of life of the elderly on Earth.

Similarly, space-flown *Caenorhabditis elegans* (1 mm length and 1,000 somatic cells) showed decreased locomotive activity and expression of muscle proteins, mitochondria-related proteins, and cytoskeletal components 7. BMP/TGF-β DBL-1 signaling and body length were reduced 7-9. Muscle cell size and strength also decreased during spaceflights 10,11. Furthermore, endogenous dopamine and *comt-4* expression levels were downregulated in *C. elegans* raised under space µG and 3D clinostat-based simulated µG conditions 12. In other words, even small nematodes exhibit pathophysiology similar to that seen in spaceflight mice and astronauts. However, these causes are unlikely to be due to simple weight loss. Recently, we found that physical stimulation intervention with small beads during culturing under simulated μG could restore locomotive activities, body length, *comt-4* gene expression, and endogenous dopamine levels 12.

Here, we studied whether stimulation intervention can mitigate nematode pathophysiology in actual space μG. Cell-autonomous gravity-sensing mechanisms cannot fully explain the coordinated responses in the tissues and organs of animals and astronauts 13. Central mechanosensory-based gravity sensing enables coordinated changes throughout the animal’s lifespan. We aimed to test the hypothesis that reduced contact stimulation in µG affects synaptic signaling. A space experiment, “Neuronal Integrated System” was conducted to investigate the role of µG in neuromuscular dysfunction and aging in *C. elegans* and whether physical stimuli could mitigate these effects in µG. In our experiment, we analyzed gene expression changes with and without physical contact stimulation using wild-type *C. elegans* and a mechanosensory-deficient *mec-4* mutant 14-16. We aligned gene expression changes with the physiological effects of µG, examining the neuromuscular impact of reduced tactile stimulation during spaceflight, including neuronal transmission attenuation and accelerated aging.

## Results

### Analysis of space µG and physical stimuli effects on gene expression in wild-type *C. elegans*

During the NIS space experiment, *C. elegans* samples were launched and cultured aboard the ISS, producing an F1 generation in space µG (Fig 1A-B). Small Bag A contained L4 larvae to day-one (D01) adults, which were frozen after the movement observation. Large Bag B was used for chemical fixation in the ISS for tissue observation. Half of the day-10 (D10) adult worms were frozen without fixation for transcriptome analysis. To restore physical contact stimulation, we placed plastic beads with a specific gravity of 1.00 g/cc and a diameter of 250-300 µm into these sample bags. The movement frequency of worms raised in space μG was slower, and the body length became shorter than those in 1G (Fig. 2A,B; Suppl Movie 1). In the bead-added condition (µGB), worms were in constant contact with beads and therefore exhibited strong bending and extension movements; however, motility could not be quantified.

**Figure 1.**
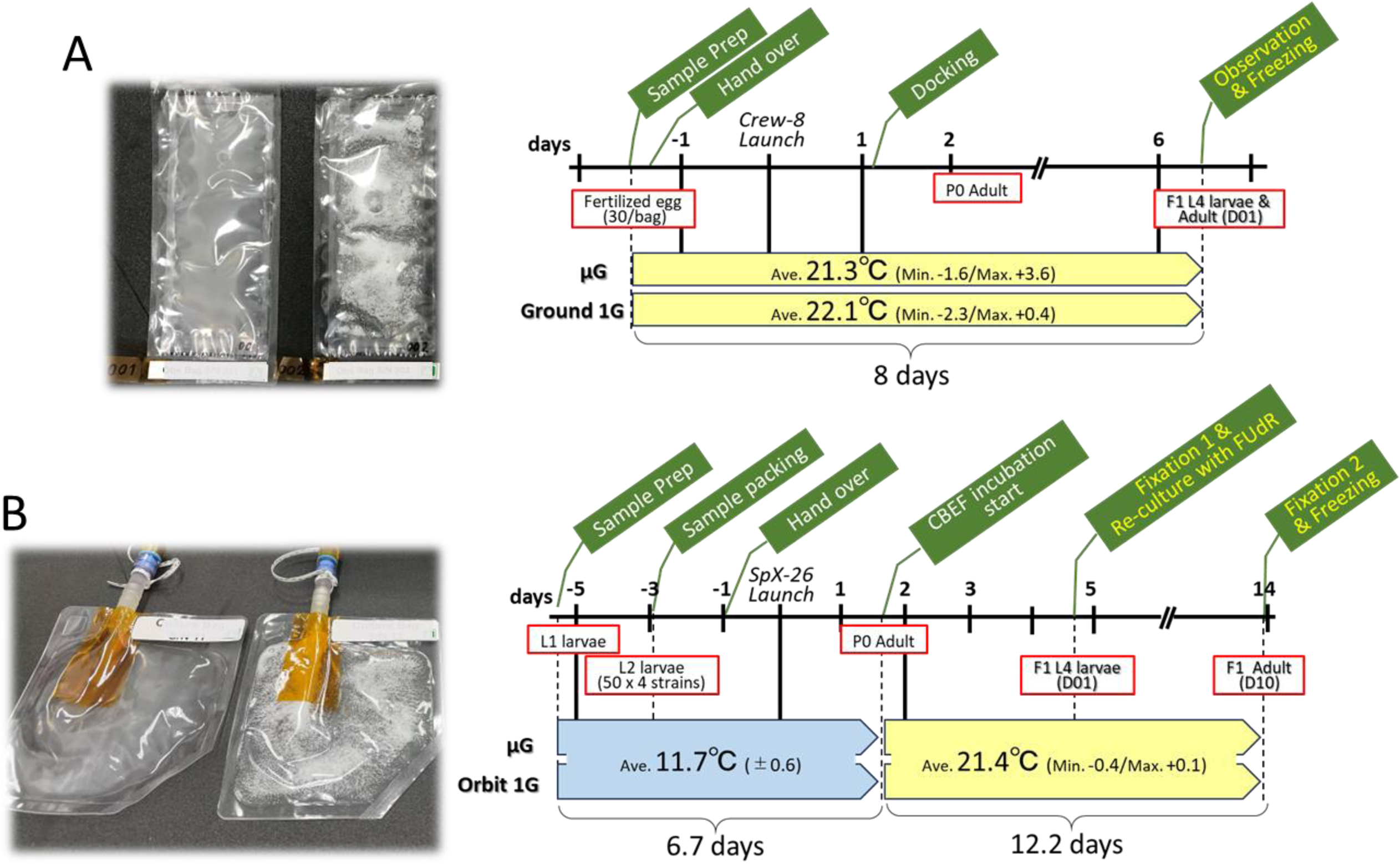
Overview of the *C. elegans* Neuronal Integrated System (NIS) experiment and cultivation status. (A) Small culture bags were used to observe F1 generation nematode motility, grown from eggs in space μG until adulthood, and to analyze gene expression after freezing. Ground 1G samples at Kennedy Space Center received identical treatment. (B) The larger bag contained 10-day-old adult worms grown in space with FudR added at D01 to inhibit the development of the next generation. Half of the samples were chemically fixed in orbit for histological analysis, while the other half were frozen for gene expression analysis to study the effects of μG aging. Orbit 1G samples underwent artificial 1G centrifugation on the ISS. The culture bags contained white plastic beads for contact stimulation.

**Figure 2.**
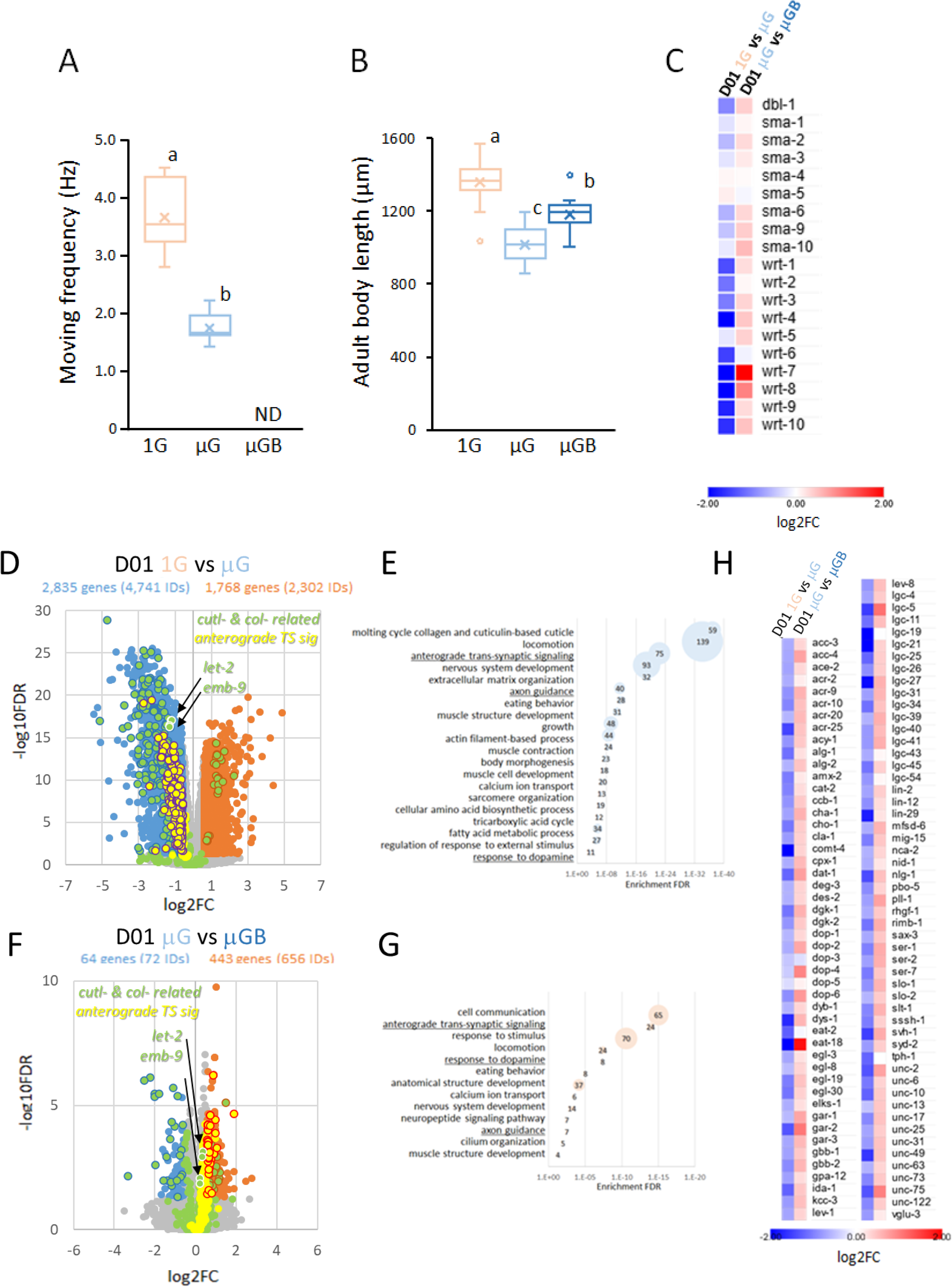
Effects of μG and bead addition under culture conditions on locomotor behavior and gene expression changes in N2 wild-type D01. (A) D01 worms’ swimming behavior frequency was calculated from video images from the JAXA Kibo module on the ISS. (B) Nematodes were cultured in bags, frozen, thawed, and the body lengths of 20 nematodes were measured. Significant differences are indicated by uppercase letters (p < 0.05, ANOVA with Tukey’s test). (C) Heatmap analysis of log2(FC) values of 1G vs. μG and μG vs. μGB expression of dbl-1, sma, and wrt genes for body length control in C. elegans. (D) Volcano plot comparing gene expression by RNA sequencing of the D01 cohort at 1G and μG. Expression ratio log2(FC) and -log10FDR were plotted using three biological replicates. Blue: decreased genes; orange: increased genes; gray: not significant; green: cuticle and collagen genes; green with white outlines: *let-2* and *emb-9*; yellow: anterograde trans-synaptic signaling and dopamine genes (purple outlines: decreased). (E) GO enrichment analysis of 2,835 genes decreased under μG vs. 1G. (F) Volcano plot comparing gene expression in the D01 cohort under μG and μGB conditions. (G) GO enrichment analysis of 443 genes increased under μGB vs. μG. (H) Heat map of 101 genes with decreased μG expression related to anterograde trans-synaptic signaling and dopamine response genes, comparing D01 1G vs. μG and μGB vs. μG expression.

However, the recovered samples showed a significant increase in body length. To investigate the molecular basis of these changes, we analyzed gene expression in the D01 cohort (L4 to young adults) grown in space µG and μGB while conducting the same analysis in a 1G environment on Earth. RNA sequencing revealed that the expression levels of certain BMP/TGF-β DBL-1 signaling genes, which are crucial for regulating body length, were reduced in the D01 cohort grown under μG conditions. Additionally, an improvement was observed in the μGB (Fig. 2C). Similar and more pronounced expression changes occurred in the hedgehog-like Warthog (*wrt*) gene family, which regulates molting and body morphology 17 (Fig. 2C).

Volcano plot analysis showed that 4,741 gene IDs (2,835 genes) were significantly downregulated (FC <2/3, FDR p<0.05), while 2,302 gene IDs (1,768 genes) were significantly upregulated (FC >1.5, FDR p<0.05) in µG (Fig. 2D: 1G vs. μG). GO enrichment analyses revealed that the downregulated genes during µG included “molting cycle collagen and cuticle-based cuticle (59/102)”, “locomotion (139/571)”, “anterograde trans-synaptic signaling (75/262)”, “nervous system development (93/427)”, “extracellular matrix organization (32/58)”, “axon guidance (40/142)”, “eating behavior (28/71)”, “muscle structure development (31/94)”, “tricarboxylic acid cycle (12/26)”, and “response to dopamine (11/35)” (Fig. 2E, Supplementary Table 1). Upregulated genes included “positive regulation of cell cycle (18/86)”, “negative regulation of cell cycle (18/99)”, “negative regulation of gene expression (39/300)”, “negative regulation of metabolic process (68/634)”, “proteasomal protein catabolic process (28/224)”, “DNA repair (37/228)”, and “histone modification (19/167)” (Supplementary Table 2).

Adding beads to µG cultures (µGB) reduced altered gene expression, with 72 gene IDs (64 genes) decreased and 656 gene IDs (443 genes) increased (Fig. 2F: μG vs. μGB). GO analysis of upregulated genes showed “anterograde trans-synaptic signaling (24/262)”, “locomotion (24/571)”, “response to dopamine (8/35)”, “nervous system development (14/427)”, “neuropeptide signaling pathway (7/140)”, and “axon guidance (7/142)” (Fig. 2G, Supplementary Table 3). The number of genes in these GO terms did not return to the levels at 1G, indicating partial recovery in the µGB.

We focused on genes (101 of 334 IDs) whose expression significantly decreased under µG related to “anterograde trans-synaptic signaling” and “response to dopamine”, including TH/*cat-2* for dopamine synthesis, COMT/*comt-4* and MAO/*amx-2* for dopamine catabolism, *dat-1* for dopamine transport, and dopamine receptor genes *dop-1* to *-5* (Fig. 2D, F). These included genes for serotonin and acetylcholine synthesis and receptors (*tph-1, cha-1, gar-1, -2, -3, ser-1, -2, and-7*). µGB increased the expression of most genes, as shown in the heatmap of 101 genes with Log2(FC) values (Fig. 2H), suggesting that neuronal signaling was altered in µG and restored by contact stimulation.

### Synaptic vesicle dynamics in the nervous system in space-grown *C. elegans*

We focused on the expression changes of 22 genes involved in neuronal vesicle trafficking, exocytosis, and synaptogenesis 18. Heat map analysis between µG vs. 1G and µG vs. µGB showed decreased expression of *liprin*/*syd-2* and *KIF1A*/*unc-104* (anterograde synaptic vesicle transport), *RIMS1*/*unc-10* (synaptic membrane exocytosis), *unc-13* (vesicle exocytosis), *cla-1* (presynaptic active zone structure), and *syg-1*/*syg-2* (synaptogenesis) in the wild-type D01 cohort in µG, and µGB restored these expressions (Fig. 3A).

**Figure 3.**
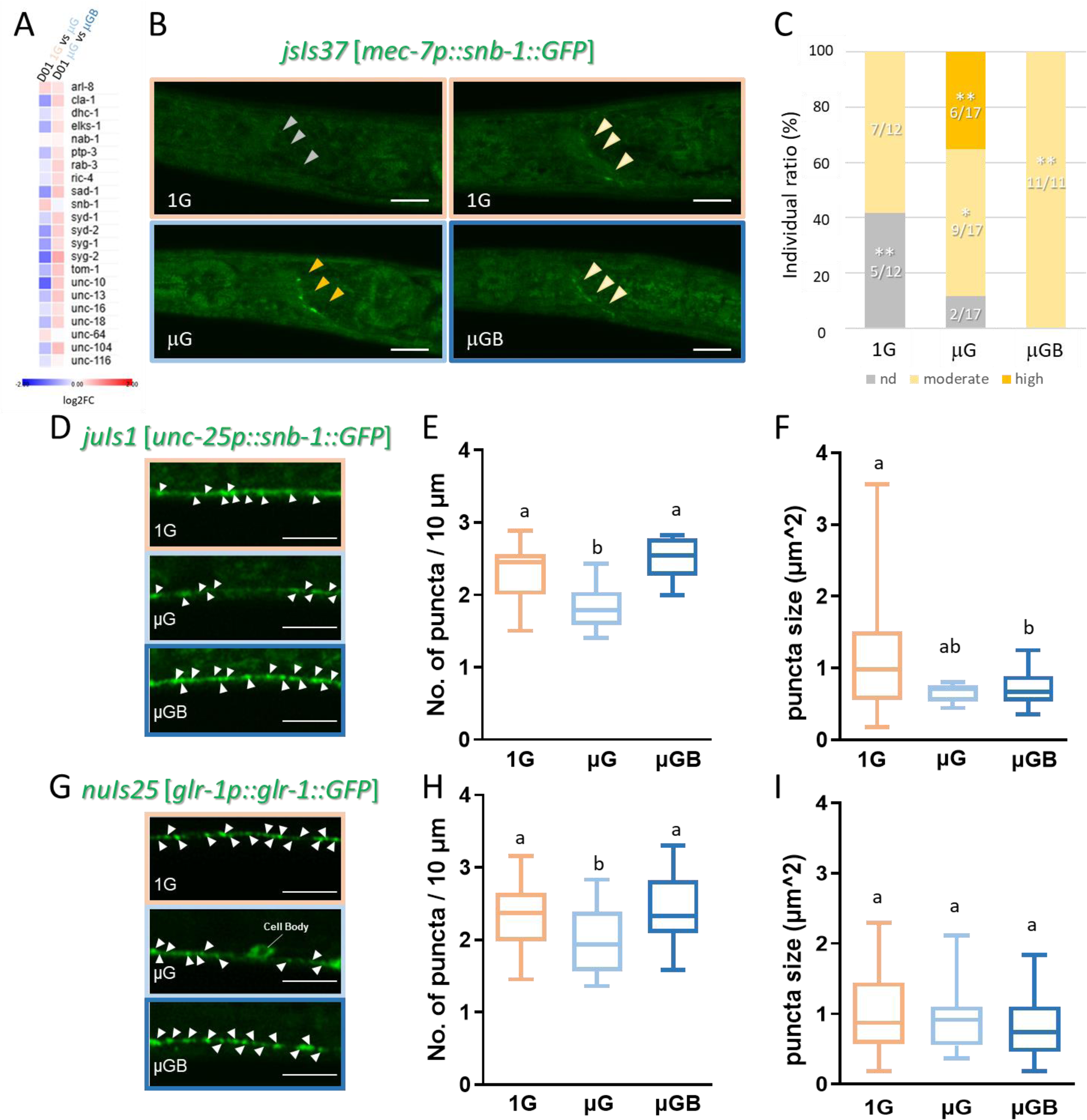
Pre-synaptic and post-synaptic fluorescent puncta under spaceflight microgravity and bead application. (A) Heatmap showing expression levels (log2FC) of synapse-related genes comparing D01 1G vs. μG and D01 μGB vs. μG from RNA sequencing (L4 to young adult cohort). (B) Images of SNB-1::GFP presynaptic puncta in ALM touch sensory neurons in the nerve ring of L4 larvae under space μG, μGB, and 1G conditions. Scale bars: 10 μm. (C) SNB-1::GFP puncta intensity in NR synapses classified as high (<1000), medium (300-1000), or not detected (<300), with proportions plotted. Sample sizes: n=12, 17, 11. Chi-square test used. *p<0.05, **p<0.01. (D) Images of SNB-1::GFP presynaptic puncta in GABAergic motor neurons of L4 larvae under μG, μGB, and 1G conditions. (E) SNB-1::GFP presynaptic puncta density per 10 μm in L4 larvae (n=20< each). (F) Synaptic puncta size (n=20< each). (G) Images of GLR-1::GFP postsynaptic puncta in L4 larvae under μG, μGB, and 1G conditions. (H) GLR-1::GFP postsynaptic puncta density (n=20< each). (I) GLR-1::GFP puncta size (n=20< each). Scale bars: 20 μm. One-way ANOVA with Tukey’s test; different letters indicate significant differences at p<0.05.

Next, we investigated synaptic dynamics in space-grown L4 larvae (Large Bag B) expressing SNB-1::GFP, a presynaptic marker that forms puncta at synapses. In worms expressing *jsIs37* [*mec-7p::snb-1::gfp*], synaptic puncta were visible in the nerve ring of ALM touch neurons (Fig 3B, 19). In space µG, 35% showed strong SNB-1::GFP signals, 53% showed moderate signals, and 12% showed faint signals. The Space 1G environment showed moderate signals in 60% of the cases and no signal in 40% of the cases, with no strong signals. With μGB stimulation, all specimens exhibited moderate signals. Space µG appears to alter synaptic vesicle release and/or accumulation in ALM neurons, whereas touch stimulation through beads suppresses these changes.

Since locomotory behavior changed in space µG (Fig 2A), we examined whether motor neuron synapse dynamics were altered. We focused on DD/VD GABAergic motoneurons that synapse with the body wall muscles controlling movement. In worms expressing *the juIs1*[*unc25p::SNB-1::GFP*] transgene 20, synaptic puncta were visible along the GABAergic motor neurons (Fig 3D). Space µG reduced presynaptic puncta density but not size compared with the space 1G control (Fig. 3D-F), indicating altered synapse development at the neuromuscular junction. Beads µGB reversed synaptic puncta density to levels seen in 1G (Fig. 3D, E).

We also investigated postsynaptic changes using GLR-1 expression. *glr1p::GLR-1::GFP* marks post-synaptic puncta in ventral nerve cord interneurons, with changes in puncta density linked to post-synapse formation defects 21.

GLR-1::GFP fluorescence changes when worms are isolated but is restored by mechanical stimulation 22. Similar to SNB-1::GFP, the density of GLR-1:: GFP puncta decreased in µG but recovered in µGB, whereas the puncta size remained unchanged (Fig. 3G-I). This suggests that space µG affects synaptic gene expression and synapse development in the nervous system, with physical stimuli ameliorating these defects.

### Gene expression changes in aged *C. elegans* in space µG (D01 vs D10)

Previous studies on *C. elegans* spaceflight have demonstrated that µG leads to muscle atrophy and hinders neuronal debris clearance, indicating accelerated aging 7,10,23,24. To study the effects of space μG and aging on differentially expressed genes (DEGs) in *C. elegans*, we performed RNA-seq analyses on D01 (a L4 to young adult cohort) and 10-day-old (D10) wild-type adults treated with FUdR. D10 worms were cultured by transferring half of the F1 generation L4 larvae to fresh culture bags with FUdR (Fig. 1B). PCA of gene expression, with biological triplicates under each condition, identified three clusters: the 1G group within the D01 cohort and the μG and μGB groups in D01 and D10 under all gravity conditions (Fig. 4A). PC1 captured age-related changes between D01 and D10, while PC2 represented growth environment changes at 1G and μG, with gravity change effects being most significant in the D01 cohort. Gene expression in the D01 μGB cohort shifted toward the D01 1G cohort along the PC2 axis.

**Figure 4.**
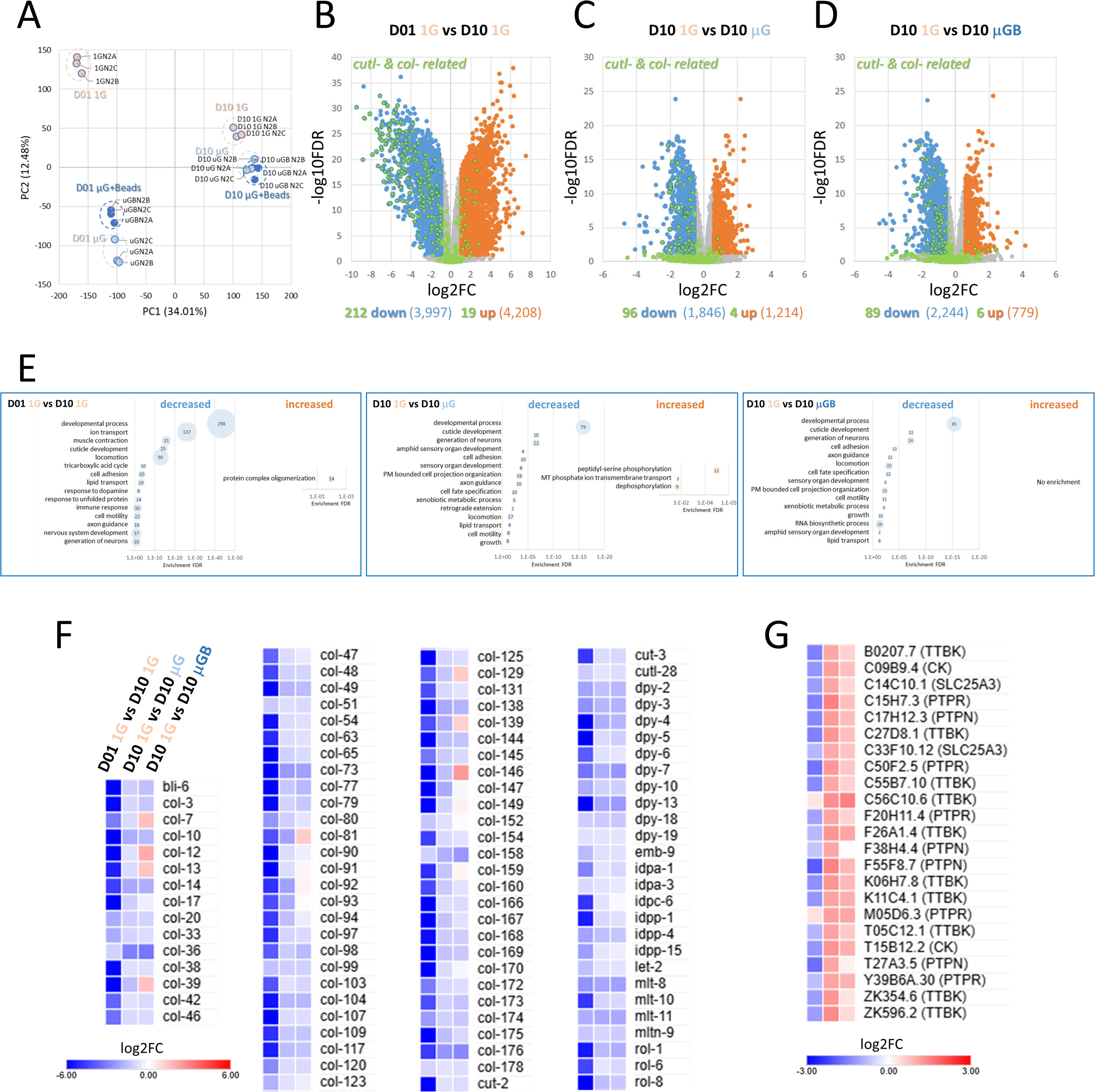
Changes in gene expression in aged worms in μG. (A) PCA analysis of gene expression with replicates in the D01 cohort (L4 to young adults) and 10-day-old (D10) adults under μG, μGB, and 1G conditions. (B) Volcano plot of age-related DEGs in D01 1G vs. D10 1G. Blue: decreased genes; orange: increased genes; gray: not significant; green: cuticle/collagen genes. (C) Volcano plot of DEGs comparing D10 1G vs. D10 μG. (D) Volcano plot of DEGs comparing D10 1G vs. D10 μGB. (E) GO enrichment analysis of DEGs from plots (B), (C), and (D). (F) Heatmap of 96 cuticle/collagen genes decreased by μG in D10 worms, comparing D01 1G vs. D10 1G and D10 1G vs. D10 μGB. (G) Heatmap of 23 TTBK and PTPs genes enhanced by μG in D10 worms, comparing D01 1G vs. D10 1G and D10 1G vs. D10 μGB.

To identify age-related genes, we examined the DEGs at D01 and D10 in the 1G environment. Volcano plot analysis showed that 3,997 gene IDs significantly decreased with age (less than two-thirds, FDR p<0.05), and 4,208 IDs increased (more than 1.5-fold, FDR p<0.05) (Fig. 4B). GO enrichment analysis revealed that genes were enriched in terms such as “developmental process”, “cuticle development”, “cell adhesion”, and neuromuscular-related terms including “muscle contraction”, “locomotion”, “response to dopamine”, and “nervous system development” (Fig. 4E, Supplementary Table 4). These enriched GO terms aligned with those downregulated in μG vs. 1G in D01 worms (Fig. 2). Of the 348 gene IDs involved in extracellular collagen, 212 showed a significant aging-related decrease under 1G conditions, while 19 showed an increase (Fig. 4B, green dot). Collagens decrease or irregularly accumulate with age, causing various aging phenomena 25,26. The increased gene IDs showed enrichment only in “oligomerization of protein complexes,” with 14 gene IDs from the TNF-alpha induced protein 1 family, containing KCTD (Fig. 4G, Supplementary Table 5). These mammalian orthologs are implicated in aging and neuropsychiatric disorders 27.

Comparing D10 aged worms under different gravity conditions, collagen gene cluster expression was more reduced in μG than in the artificial 1G environment of the ISS (Fig. 4C; D10 1G vs. D10 μG). GO analysis of DEGs with decreased expression in D10 μG showed enrichment of neuron-related terms, “locomotion” and “cuticle development” (Fig. 4E middle panel; Supplementary Table 6). Genes highly expressed in D10 μG showed enrichment of GO terms related to “peptidyl-serine phosphorylation” including tau-tubulin kinase (TTBK) family, “mitochondria phosphate ion transmembrane transport” and “dephosphorylation” involved in the protein tyrosine phosphatase (PTPs) family (Fig. 4C, E; Supplementary Table 7). Mammalian TTBKs are implicated in neurodegenerative diseases and aging, with their overexpression inducing axonal degeneration 28.

When comparing D01 1G vs. D10 1G, as well as D10 1G vs. D10 μG, and D10 1G vs. D10 μG B, it was found that among the 96 gene clusters related to the cuticle and collagen, whose expression decreased with aging under 1G, expression declined further under μG, while the decrease was somewhat alleviated by μGB (Fig. 4D; Supplementary Table 8). In the D10 μGB group, eight collagen genes were notably upregulated (Fig. 4F). Conversely, in D10 μGB, the increased expression of 23 TTBK and PTPs family genes in D10 μG was suppressed, and no significant enrichment of increased GO terms was observed (Fig. 4G, E).

### Age-related changes in neurons under μG conditions

We examined the effects of μG on synapse density and size in D10 aged worms. The density of SNB-1::GFP presynaptic puncta in DD/VD motor neurons showed no difference between D10 μG and D10 μGB when compared to D10 1G, but it was reduced in D10 μG compared to that in D10 μGB (Fig. 5A, B). Conversely, the puncta size increased in the D10 μG group and was suppressed in the D10 μGB group (Fig. 5C). Analysis of GLR-1::GFP post-synaptic puncta showed decreased density in the μG group compared to the 1G controls, with beads restoring the density (Fig. 5D, E).

**Figure 5.**
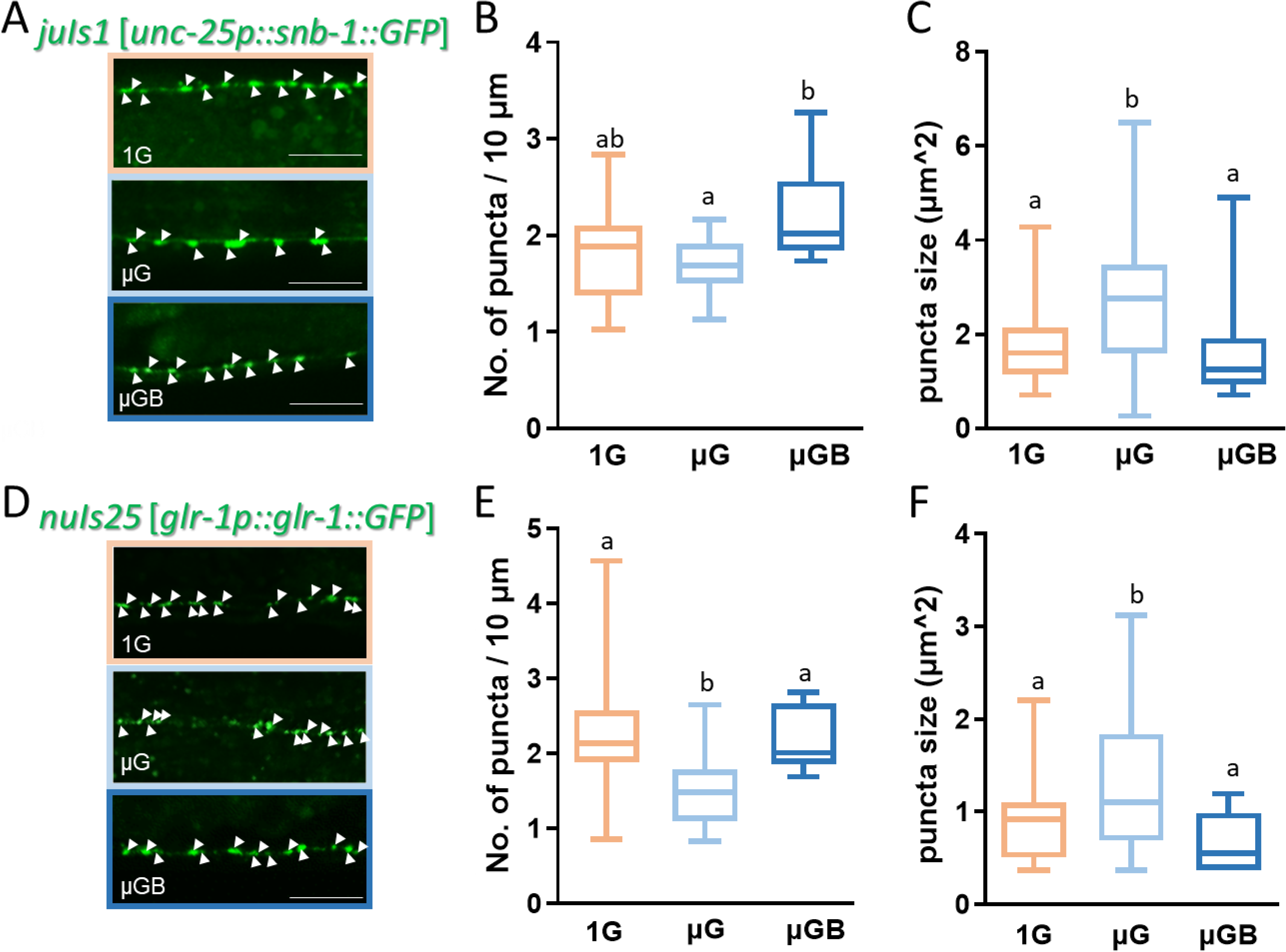
Age-associated synaptic changes in SNB-1::GFP and GLR-1::GFP under space μG and μGB conditions. (A) SNB-1::GFP presynaptic puncta images in GABAergic motor neurons of D10 aged worms under space 1G, μG, and μGB conditions. (B) SNB-1::GFP presynaptic puncta density calculated in D10 aged worms (n = 20< each). (C) Synaptic puncta size calculated in D10 aged worms (n = 20< each). (D) GLR-1::GFP postsynaptic puncta images in D10 aged worms under space 1G, μG, and μGB conditions. (E) GLR-1::GFP postsynaptic puncta density calculated in D10 aged worms (n = 20< each). (F) GLR-1::GFP postsynaptic puncta size was calculated (n = 20< each). Scale bars: 20 μm. Statistical analysis was performed using one-way ANOVA with Tukey’s test; shared letters indicate no significant difference (p < 0.05).

Worms aged in μG exhibited an increased puncta area, but the addition of beads restored this to 1G levels (Fig. 5F). The size and intensity of these puncta correlate with vesicle accumulation and or decreased release 29,30.

To examine whether space μG accelerates neuronal aging, we monitored DD/VD motor neuron axon degeneration using *unc25p::GFP*. In the ground-based experiment conducted at 1G, we noted axonal defects such as branching, truncation, and process merging by D10, with significant blebs forming along the axons by D15 (Suppl. Fig. 1). In L4 larvae (D01) raised in space, the proportion of individuals with no damage in the µGB remained unchanged, while there was an increase in the proportion of individuals exhibiting a slight rise in blebs (11-15 per individual) (Fig. 6). In D10 aged worms, µG increased both blebs and axonal defects compared to the 1G space control. Bead-loaded D10 µGB restored the number of blebs and axonal defects to the levels observed in the 1G control, suggesting that physical stimulation can mitigate neuronal damage accelerated by µG aging.

**Figure 6.**
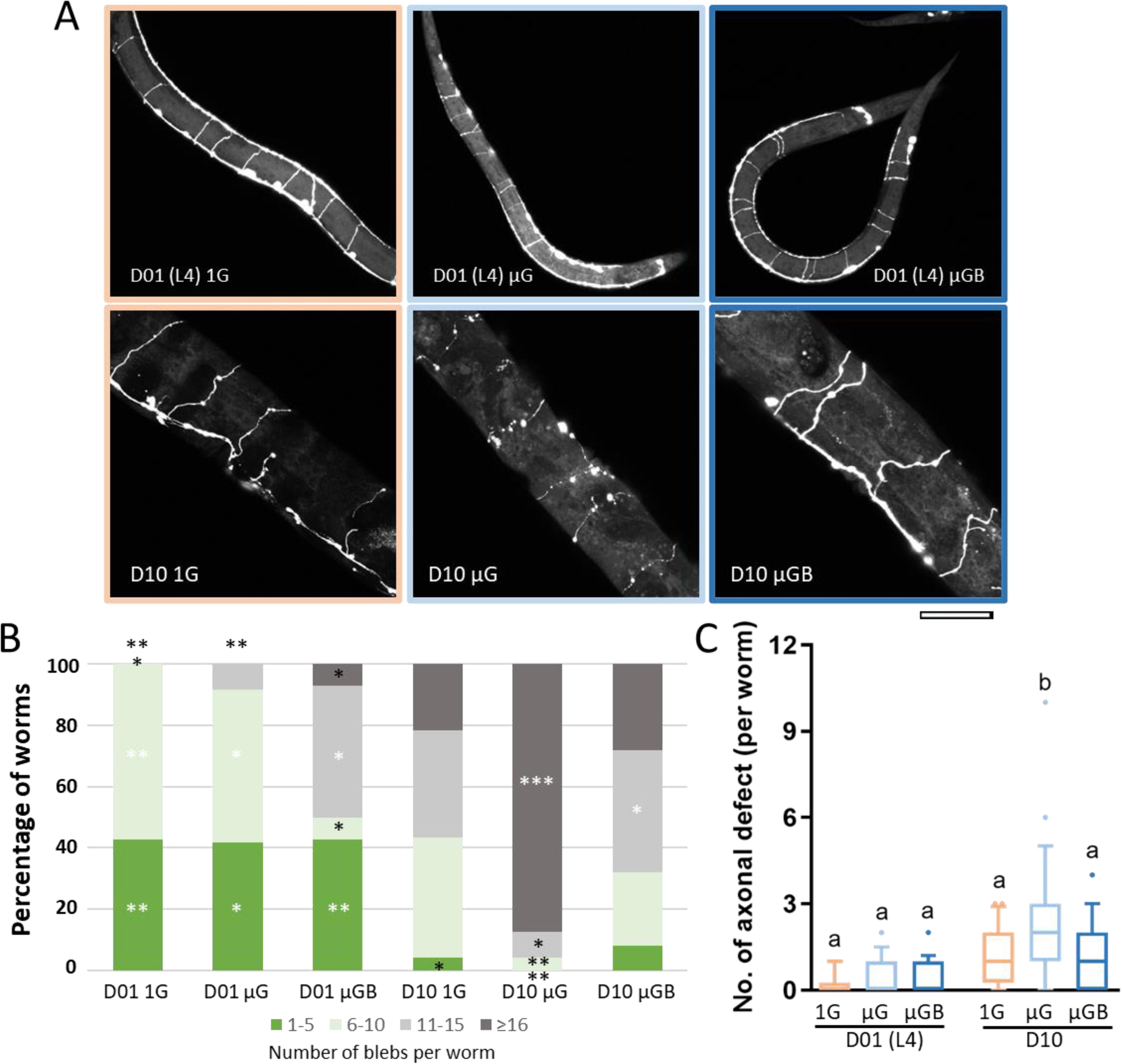
Changes in GABAergic motor neuron morphology during aging in space μG. (A) Images of SNB-1::GFP-labeled axon commissures of DD/VD GABAergic motor neurons (*juIs76* [*unc-25p::SNB-1::GFP*]) near the vulva of L4 larvae (D01) and D10 aged worms under space μG, μGB, and artificial 1G conditions. Scale bar: 50 µm. (B) Blebs in four axon commissures were categorized by damage level, and individual proportions were plotted on a stacked graph. Sample sizes from left to right: n=14, 12, 14, 23, 24, and 25. Significance was tested using a 6×4 chi-square test. *p<0.05, **p<0.01, ***p<0.001. (C) Analysis of axonal defects, including branching, severing at the commissures, and improper rotation and fusion along the axon. The same individuals as in (B). Statistical significance was determined using one-way ANOVA with Tukey’s post-hoc test. Different letters indicate significant differences at p<0.05.

Using *the dat-1p::GFP* expression strain, we examined age-related changes in dopaminergic sensory neuron CEPs dendrites. In D10 aged adults, dendritic damage in CEPs was less severe than DD/VD motor neuron axonal dysfunction. The D10 μG group exhibited a slight increase in age-related damage that was not mitigated by µGB (Suppl Fig. 2).

### Acceleration of muscle mitochondria senescence in space μG

*C. elegans* body wall muscle cells (BWMCs) contain sarcomeres and mitochondria in an observable monolayer. With aging, the mitochondrial network structure fragments and volume decrease, while mitochondrial Ca^2+^ increases before sarcomere collapse 31-33. Using transgenes *ccIs4251* (*mitoGFP*, *nucGFP*) and *aceIs1* (*mito LAR-GECO*), we found that mitochondrial networks in L4 larvae (D01) remained unchanged in both 1G and μG environments with or without beads (Suppl. Fig. 3). However, D10 μG-aged adults showed significant mitochondrial fragmentation, swelling, and volume reduction (Fig. 7A, B, Suppl. Fig. 4). Over 50% of BWMCs showed severely damaged mitochondria, while space 1G conditions showed only 15% severe damage. Furthermore, mitochondrial Ca^2+^ levels were higher in D10 μG worms than in space 1G controls (Fig. 7C). The addition of beads (D10 μGB) suppressed accelerated muscle mitochondrial aging, returning the levels near those of the D10 1G control (Fig. 7B, C). In contrast, myofibers remained almost intact, while mitochondrial aging accelerated in D10 μG (Fig. 7A).

**Figure 7.**
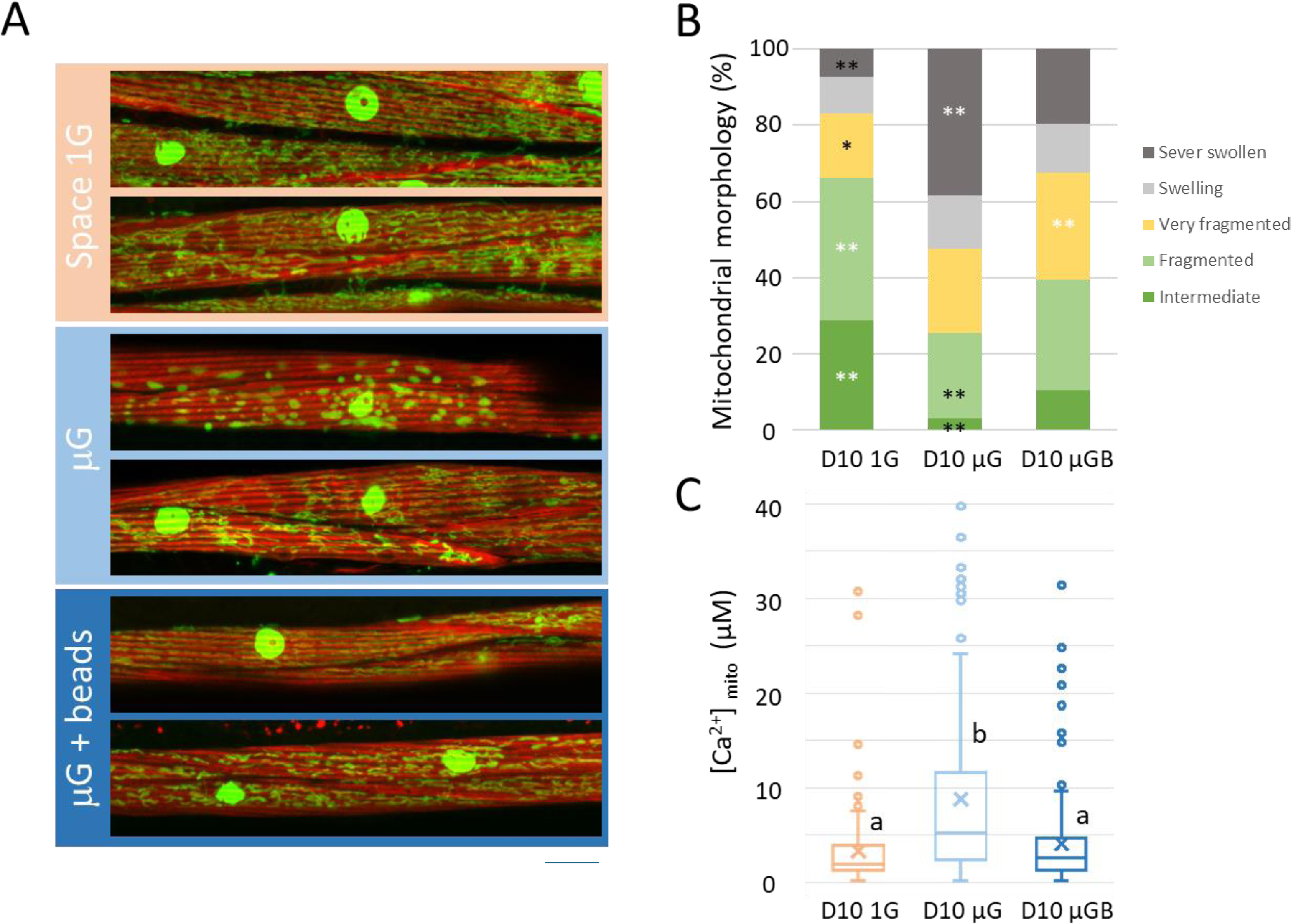
Muscle mitochondria in aged worms in space μG. (A) Images of body wall muscle cells from D10 nematode ATU3301 cultured under space μG, μGB, and artificial 1G conditions. Nuclear and mitochondrial GFP and muscle actin filaments were visualized using rhodamine phalloidin. Scale bar: 10 μm. (B) Analysis of muscle cells with abnormal mitochondrial morphology (n = 250-300 cells from 40+ worms). Mitochondrial changes were classified as “intermediate”, “fragmented”, “very fragmented”, “swelling”, and “severely swollen” as shown in Supplementary Figure 4. The chi-square test was used to determine significance. *p<0.05, **p<0.01, ***p<0.001. (C) Analysis of mitochondrial Ca^2+^ concentrations ([Ca^2+^]_mito_) in muscle cells using mtGECO intensity. [Ca^2+^]_mito_ was calculated as described in 32. Significance was determined using one-way ANOVA with Tukey’s test. Different letters indicate significant differences at p<0.05.

### Effect of mechanoreceptors on body-size and global gene expression changes in space μG

We demonstrated that restoring physical stimulation with beads can alleviate the effects of space µG in *C. elegans*, suggesting that the lack of tactile stimulation may drive μG perception. To assess the role of touch sensation in μG signaling, we examined changes in body length in N2 wild-type, *mec-4* (*u253*), *trp-4* (*ok1605*), and *cat-2* (*e1112*) mutant strains. The *cat-2* (*e1112*) mutants, which are defective in dopamine biosynthesis and are normally longer than wild-type 34, showed a reduced body length under μG, similar to wild-type, and the addition of beads improved their body length (Fig. 8A). This suggests that dopamine depletion under microgravity in space 12 is not the cause of μG-dependent body length shortening. The *trp-4* (*ok1605*) mutant, involved in both proprioception and posterior harsh touch 35, showed reduced length under μG, with a further reduction when beads were added. However, *mec-4* (*u253*) mutants showed no length difference between the 1G and μG environments, and bead addition had no effect (Fig. 8A). These results indicate that the μG-induced decrease in body length is mainly mediated by MEC-4 mechanoreceptors, whereas the effect of bead addition under μG may involve TRP-4 mechanoreceptors.

**Figure 8.**
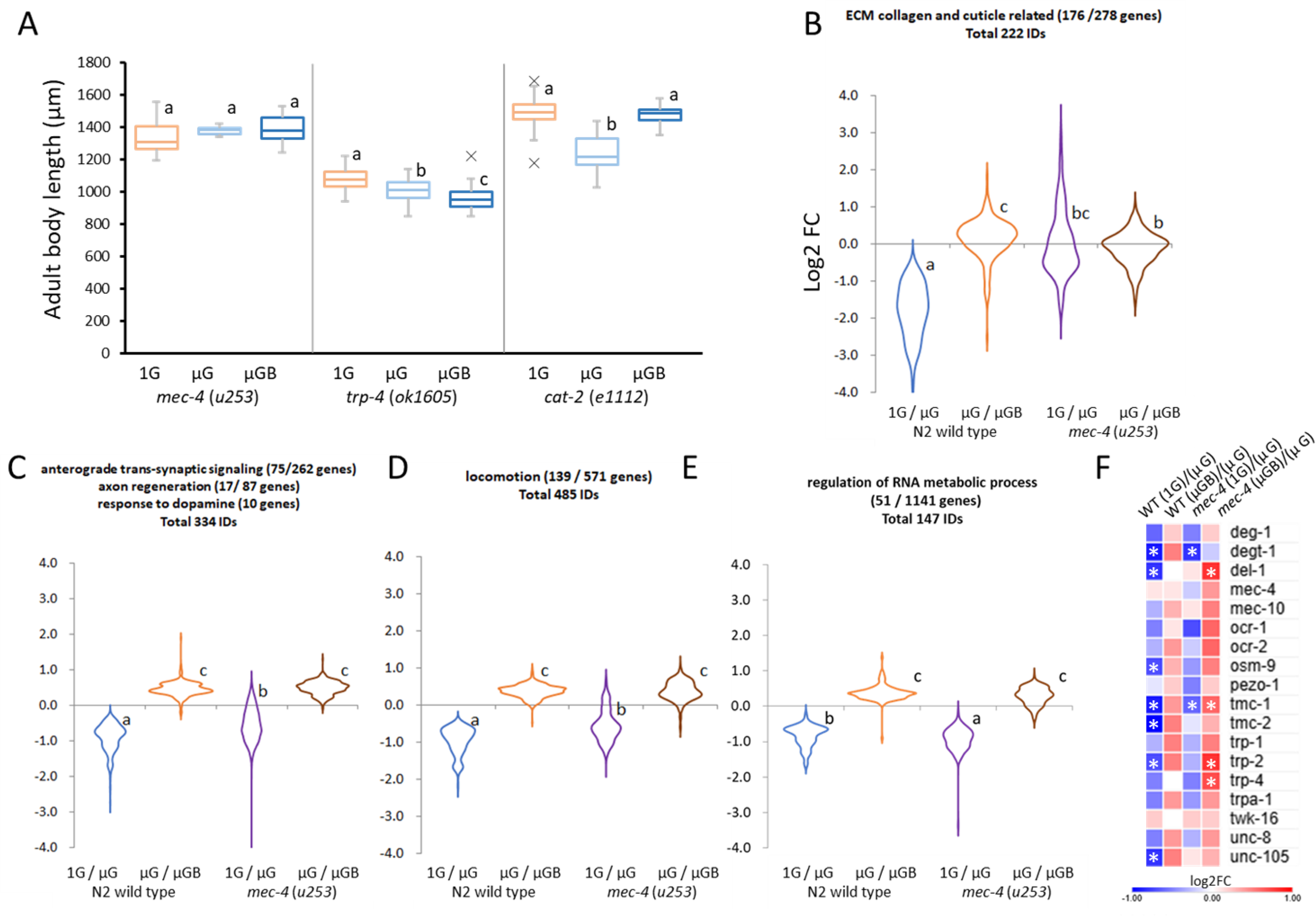
Body length and gene expression changes in mutants of mechanoreceptor genes grown under space μG, μGB, and terrestrial 1G. (A) D01 *mec-4*, *trp-4*, and *cat-2* mutants were cultured, frozen, thawed, and the top 20 nematode body lengths were measured. Different uppercase letters indicate significant differences (p < 0.05, ANOVA with Tukey’s test). (B-D) Gene expression changes of “ECM collagen and cuticle-related” GO genes (B: 222 IDs), “anterograde trans-synaptic signaling,” “axon regeneration,” “response to dopamine” GO genes (C: 334 IDs), “motility” GO genes (D: 485 IDs), and “RNA metabolic processes” GO genes (E: 147 IDs) were compared between wild-type and *mec-4* mutants using log2(FC) ratios of 1G vs. μG and μG vs. μGB. (F) Heatmap analysis of the expression ratios of 18 mechanoreceptor genes between D01 wild type and *mec-4* mutants under different gravity conditions. * indicates genes with <0.67 or >1.5-fold expression at FDR p<0.05.

Gene expression analyses were performed on *mec-4* (*u253*) mutants (D01 L4 to young adult cohort) raised under μG and μGB conditions and compared with those raised in a 1G environment. In the μG vs. 1G comparison, 176 ECM collagen- and cuticle-related genes that were downregulated in D01 wild-type remained unchanged in *mec-4* (*u253*) mutants, with some genes upregulated (Fig. 8B, Suppl. Table 9).

Expression of “anterograde trans-synaptic signaling” genes (334 IDs) and “locomotion” genes (485 IDs) in *mec-4* (*u253*) mutants were reduced under μG, but less than in wild-type (Fig. 8C, D, Suppl. Table 9). Expression of 51 genes involved in “regulation of RNA metabolic process” was downregulated by μG in both wild type and *mec-4*.

With beads (μGB), the expression of these three GO term genes returned to 1G levels in both strains (Fig. 8C-E, Suppl. Table 9), whereas collagen- and cuticle-related genes showed no change in *mec-4* (*u253*) mutants. The findings indicate that the reduction in body length under μG is primarily associated with decreased expression of ECM collagen and cuticle-related genes due to weakened MEC-4 mechanoreceptor tactile signaling. Additionally, there were moderate MEC-4-mediated alterations in synaptic signaling and locomotion genes and MEC-4-independent changes in RNA metabolic processes.

We wondered whether other mechanoreceptors may mediate genetic and physiological changes under µG. We evaluated the expression changes of 18 genes encoding major mechanoreceptors 36, comparing wild type and *mec-4* (*u253*) under μG conditions, with and without beads. The expression of seven mechanoreceptor genes was significantly reduced (less than two-thirds, FDR p<0.05) under µG conditions in the D01 wild-type cohort (Fig. 8F marked with *), with most of the remaining genes showing reduced expression, except *mec-4* and *twk-16*. This suggests that the absence of tactile stimuli downregulates most mechanoreceptors. Two genes, *degt-1* and *tmc-1*, showed reduced expression in both wild-type and *mec-4* (*u253*) mutants in µG, indicating their independence from MEC-4. The expression of *unc-105* and *del-1* was reduced only in the wild type, suggesting MEC-4-mediated µG-dependent suppression. Suppression of *omc-9*, *tmc-2*, and *trp-2* was marked in wild-type but less in *mec-4* (*u253*), while *ocr-1* showed the opposite suppression. Bead-loaded μGB restored transcriptional repression of most mechanoreceptor genes in both strains, with greater improvement in the *mec-4* (*u253*) mutant (Fig. 8F).

### Impact of the *mec-4* (*u253*) mutation on neuromuscular aging in a 1G terrestrial environment

We studied DEGs by comparing the D01 wild-type and *mec-4* (*u253*) cohorts (L4 to young adult) at 1G control. GO analyses showed that downregulated genes were enriched in “molting cycle collagen and cuticulin-based cuticle”, “ion transport”, and neuromuscular terms such as “sensory organ morphogenesis” and “neuropeptide signaling pathway” (Suppl. Fig. 5). These GO terms partially aligned with the genes downregulated in 1G vs. μG in the D01 wild-type cohort (Fig. 2D, Suppl. Fig. 4). The upregulated genes were enriched in “reproduction”, “autophagy”, “autophagy of mitochondrion”, “aggrephagy”, “microtubule organization”, and “nervous system development” (Suppl. Fig. 5). Twenty-one autophagy and aggrephagy genes were upregulated in *mec-4* mutants compared to those in the wild type (Fig. 9A), similar to the comparisons between spaceflight and terrestrial aging in wild type (Fig. 9A). These results suggest that decreased touch stimuli via MEC-4 at 1G may accelerate aging-related neuromuscular changes, similar to those observed under μG conditions.

**Figure 9.**
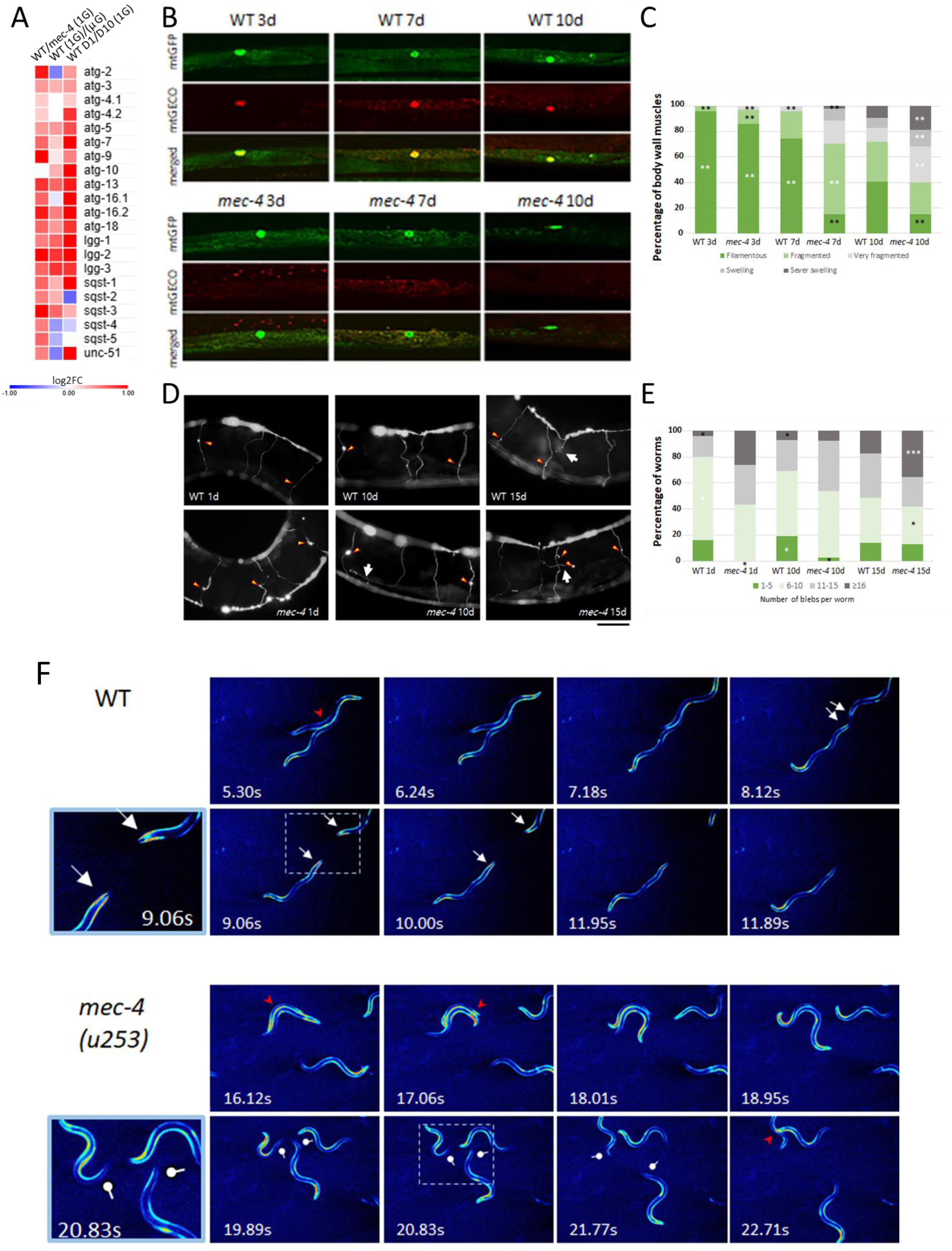
Effect of *mec-4* mutations on neuromuscular aging and muscular calcium transients in a terrestrial 1G environment. (A) Heatmap analysis of log2(FC) ratio of autophagy/aggrephagy genes between D01 wild type and *mec-4* mutants in 1G environment, plotted between N2 wild-type D01 1G vs. μG and D01 vs. D10 in 1G. (B) Images of muscle mitochondrial aging in *mec-4* mutants vs. wild type in 1G culture. Scale bar: 10 μm. (C) Mitochondrial changes were classified as shown in Supplementary Figure 4, using chi-square test analysis (n = 75–250 muscle cells from 15+ worms each). *p<0.05, **p<0.01. (D) Images of age-related changes in SNB-1::GFP-labeled axon commissures of GABAergic motor neurons in wild-type and mec-4 mutants in 1G culture. Scale bar: 50 µm. (E) Blebs in four axon commissures were categorized according to the damage level and plotted (n = 25, 23, 42, 39, 35, and 31). Chi-square test: *p<0.05, **p<0.01, ***p<0.001. (F) Cytosolic calcium transients during crawling and contact stimulation in wild-type and *mec-4* (*u253*) adult mutants cultured on *E. coli* OP-50 NGM plate. Calcium concentration changes were visualized using *goeIs3* [*myo-3p::SL1::GCamP3.35::SL2::unc54 3’UTR* + *unc-119*(*+*)] and converted using Royal Color. Sequential clips at 1-second intervals were shown in the videos (Supplementary Movie 2).

We compared muscular mitochondrial collapse and DD/VD motor neuron axon degeneration between wild-type and *mec-4* (*u253*) mutants. Analysis revealed enhanced mitochondrial aging, including fragmentation and swelling. Smaller punctate mitochondria showed higher Ca^2+^ accumulation, suggesting the potential occurrence of mitophagy 32 in *mec-4* (*u253*) mutants (Fig. 9B, C). Similar to muscle mitochondrial impairment, axonal blebbing of DD/VD motor neurons occurred earlier in *mec-4* (*u253*) mutants than in wild-type (Fig. 9D, E). However, under space μG conditions, wild-type D10 exhibited more pronounced muscle and motor neuron aging damage than *mec-4* (*u253*) mutants at terrestrial 1G, indicating that the effects of space μG extend beyond merely reducing MEC-4 channel stimulation.

The *mec-4* (*u253*) mutant lacks mechanostimuli-activated calcium transients in ALM touch neurons during gentle touch stimuli 16. We examined whether *mec-4* (*u253*) affected muscle cytoplasmic calcium transients during contact stimulation compared to wild type under 1G conditions, using *goeIs3* and *aceIs1* transgenes. No differences were observed between wild-type and *mec-4* (*u253*) mutants in calcium transients within the inner contracted myocytoplasm during crawling on agar medium (Fig. 9F, Suppl. movie 2). The calcium transients during vigorous contact between the nematodes were also similar (Fig. 9F red arrowheads; Suppl. movie 2). However, after weak friction when worms meet and separate, calcium transients occur in the tail muscle cytoplasm of wild type but not *mec-4* (*u253*) mutants (Fig. 9F white arrows, circular heads; Suppl. movie 2). These results showed *that the mec-4* (*u253*) mutation eliminated calcium transients in body wall muscle cells upon weak tactile stimulation and ALM touch neurons.

## Discussion

### Global gene expression changes with decreased contact stimuli in space μG

In this study, we confirmed the downregulation of specific genes associated with the muscle, cytoskeleton, mitochondria, TGF-β/BMP *dbl-1*, and *comtt-4* in *C. elegans* raised in space μG, consistent with findings from previous research 7-12. We also observed a decrease in the expression of genes related to anterograde trans-synaptic signaling and dopamine response, including those involving acetylcholine and serotonin neurotransmitters, as well as genes associated with cuticle and collagen, particularly within the hedgehog-like Warthog (*wrt*) gene family in the F1 generation of the D01 cohort (L4 larvae to young adults) in space μG (Figs. 2,3,8). In comparing gene expression changes between *mec-4* (*u253*) loss-of-function mutants and wild type, it was found that the reduced expression of ECM collagen and cuticle-related genes under μG was primarily due to diminished MEC-4 mechanosensory signaling (Fig. 8). In contrast, MEC-4 moderately reduced synaptic signaling and genes associated with locomotion, whereas RNA metabolic processes were reduced independently.

Notably, reductions in mechanosensor channel expression induced by μG were also observed, whether they were dependent on, moderately linked to, or independent of MEC-4 function (Fig. 8F). Seven of the 18 mechanoreceptor genes were significantly reduced under µG conditions in D01 N2 wild type. Suppression of *unc-105* and *del-1* occurred in an MEC-4 dependent manner. Both are DEG/ENaC channels, with DEL-1 expressed in the stretch-sensitive regions of motor neurons 37, while UNC-105 functions as a stretch-sensitive channel in body wall muscle cells 38. UNC-105 also interacts with LET-2 type IV collagen, which forms a heterotrimer with another type IV collagen, EMB-9, and plays a significant role in mechanotransduction and aging 26,39. The expression of these type IV collagen genes decreased with age and was further diminished in space μG at D01 and D10 compared to the 1G environment (Figs. 2D, 4F). Furthermore, other mechanoreceptor genes*, degt-1* and *tmc-1*, which were also suppressed in μG but are independent of MEC-4, play roles in regulating foregut motility 40 and in anti-aging processes across neurons 41, respectively.

Recent research has shown that the mechanoreceptor DEGT-1 acts as a proprioceptor and localizes to the neuronal soma of pharyngeal (the foregut, equivalent to the “stomach”) neurons, where it contacts the pharyngeal basement membrane 40.

Downregulation of *degt-1* results in an increased pharyngeal pumping rate, potentially allowing adaptation to changes in microbial food conditions and the pharyngeal environment by reducing the shear force under μG. Similarly, in humans under μG conditions, the body cannot rely on gravity to move stomach contents into the intestine and must rely on peristalsis (rhythmic muscle contractions), making “forces from ingested material” unreliable sensory cues.

TMC-1 is involved in alkaline pH avoidance and mechanoperception, while studies show its neuroprotective role occurs through the TMC-1–GABA - PLCβ–DAG - PKC signaling pathway 41. Under μG conditions, the genes in this pathway—*tmc-1*, *unc-13*, *unc-31*, *gbb-1*, *gbb-2*, *egl-30*, *egl-8* (*PLCβ*), and *dgk-1*—exhibited reduced expression (Figs. 2, 3, 8). The downregulation of TMC-1 signaling may be associated with more severe damage to DD/VD motor neurons under μG conditions than a single *mec-4* (*u253*) mutant on the ground. These findings indicate that reduced tactile stimulation in space affects both MEC-4 and other mechanoreceptors, such as those involved in the stretch and proprioceptive responses of the motor organs. This may potentially initiate a negative cycle in which the expression of these receptors is suppressed, leading to dysfunction in the tissues where they are expressed and accelerating senescence.

### Neural integration changes in space μG development

We observed not only repression of anterograde trans-synaptic signaling, dopamine response, and synaptic gene expression, but also accumulation of synaptic vesicles in the nerve ring, changes in pre- and postsynaptic vesicle dynamics, and suppression of motor behavior in worms raised in space μG (Figs. 2, 3). Contact stimuli in space (μGB) restored gene expression, synaptic dynamics and motility.

In *C. elegans*, activity-dependent synaptic changes are mediated by transcription factors controlling synaptic genes, including *cla-1*, *elks-1*, *syd-2*, and *unc-10*, which are repressed by space μG (Fig. 3A). Increased neuronal activity enhances synaptic gene expression and puncta intensity, accelerating behavioral responses 42. One hypothesis suggests that μG and tactile stimulus loss may decrease motor neuron activity, leading to synaptic gene repression and altered synaptic dynamics. A study found that nematodes raised in isolation on standard agar medium showed decreased responsiveness to mechanical stimulation and shortened body length 43. Mechanical tap stimuli during the L3 larval stage can restore both reduced responsiveness and shortened body length in adults. In experiments with *mec-4* (*e1611*) mutants, which cause touch neuron degeneration and loss of touch sensitivity 14, no differences in body length were observed between isolated and grouped worms. This aligns with the physiological changes observed in worms in μG environments.

In the brains of mice exposed to long-term spaceflight, a reduction in dopamine-related gene expression was observed, but not in mice subjected to tail suspension 5,6. Video recordings of mice during spaceflight showed that, despite μG conditions, they maintained normal eating, sleeping, and defecation patterns 44.

However, there was decreased contact between the limbs/hairy skin and the culture box surfaces. C tactile fibers in the foot and hairy skin sense light touch, slow pain, and temperature and secrete oxytocin upon sensory stimulation 45. The interaction between oxytocin and dopamine influences maternal behavior and infant brain development 46,47. Infant massage stimulates growth factors, oxytocin, opioids, and dopamine through C-tactile and Aβ fibers, contributing to infant neurological development. These findings suggest that metazoan growth under μG conditions, where tactile stimulation is reduced, may significantly threaten neuromuscular development.

### Age-dependent neuromuscular changes in space

Space μG exacerbated the age-related downregulation of cuticle- and collagen-related genes compared to D10 in 1G (Fig. 4C, F). Teuscher et al. (2024) 48 found that cuticle collagen (col) mRNA levels decreased with age, and protein levels declined in the ECM. They reported that the overexpression of *col-10*, *-13*, or *-120* extended the lifespan of *C. elegans*. In addition, longevity interventions and mechanical loading increased the expression of collagen and ECM remodeling enzymes, suppressed collagen crosslinks, prevented ECM-cell detachment, and formed a regulatory feedback loop through mechanosensitive hemidesmosomes. In our space experiments, adding beads under μG conditions suppressed space μG-induced aging of synapses, motor neurons, and body wall muscles (Figs. 5-7), while upregulating eight collagen genes, including col-13 (Fig. 4F).

Beyond changes in the extracellular matrix, mitochondrial alterations in C. elegans BWMCs are associated with aging, longevity and maximum velocity locomotion 31-33,49. Our previous report 32 identified that age-related mitochondrial fragmentation, volume reduction, and swelling are driven by excessive mitochondrial Ca^2+^ accumulation and mitophagy. D10 worms raised in space μG showed enhanced mitochondrial aging, which was improved by adding beads, bringing acceleration levels closer to those under artificial 1G on orbit. Gene expression analysis revealed increased autophagy-related genes between D01 and D10, with genes upregulated at D01 under μG conditions and in *mec-4* (*u253*) mutant compared to the wild type. These results indicate accelerated muscle mitochondrial aging characteristics in C. elegans grown under reduced physical stimuli in μG.

In aging D10 worms cultured in μG, accelerated pre- and post-synapse destruction, axonal damage in DD/VD motor neurons, and mitochondrial changes were observed (Figs. 5-7). Laranjeiro et al. (2021) 23 reported that worms spending 5 days as adults on ISS showed hyperbranching in PVD and tactile receptor neurons, with accumulated neuronal waste products. This evidence suggests that spaceflight may induce premature neuromuscular tissue aging, raising concerns about the effects of long-term missions on astronauts’ health and life expectancy.

In conclusion, the μG unloading environment in space leads to rapid atrophy of bones and muscles, as observed in both spaceflight mice and human astronauts 1,2. NASA’s GeneLab data underscore mitochondrial stress as a central factor during spaceflight 50. In cultured cells, μG quickly remodels the cytoskeleton and focal adhesions 1,51. Moreover, in *C. elegans*, which consists of only 1,000 somatic cells and serves as an intermediate model between individual and cellular levels in mammals, we demonstrated that the primary effect of space μG is due to the reduction in tactile stimulation associated with levitation. This results in reduced neuromuscular integrity and accelerated aging in space; however, restoring tactile input helps alleviate these pathophysiological conditions. This finding could be crucial for sustaining homeostasis as humans continue to inhabit and evolve in space over prolonged periods.

## Materials and methods

### C. elegans strains

Wild-type N2 Bristol was used for the moving behavior analysis. Gene expression analyses used N2 and TU253 *mec-4* (*u253*) strains, while body length measurements used N2, TU253, VC1141 *trp-4* (*ok1605*), and CB1112 *cat-2* (*e1112*) mutants. NM664 *jsIs37*, TG2435 *vtIs1*, CZ13799 *juIs76*, CZ333 *juIs1*, and KP1148 *nuIs25* were used to visualize neurons and synapses. ATU3301 *ccIs4251*; *aceIs1* visualizes mitochondrial morphology and calcium in body wall muscle cells. ATU2301 *goeIs3*; *aceIs1* visualized cytoplasmic and mitochondrial calcium. ATU3301, ATU2301, TG2435, and CZ13799 were crossed with *mec-4* (*u253*) to create ATU3311 *mec-4* (*u253*); *ccIs4251*; *aceIs1*, ATU2311 *mec-4* (*u253*); *goeIs3*; *aceIs1*, YUW145 *mec-4* (*u253*); *vtIs1*, and YUW146 *mec-4* (*u253*); *juIs76*. Details about all mutant alleles and transgenes can be found in WormBase (https://www.wormbase.org/#012-34-5).

### Spaceflight experiment

The mission was conducted by the Japan Aerospace Exploration Agency (JAXA). All strains were cultured for three generations at the Kennedy Space Center Laboratory (Orlando, Florida USA,) to achieve synchronized growth. Culture bags (Fig. 1A) were filled with 8 ml of *E. coli* OP-50 solution (OD600=2.) in M9 buffer with cholesterol (M9C: KH2PO4 3 g, Na2HPO4 12H2O 15.1 g, NaCl 5 g, 1 mL of 1 M MgSO4, and 1 mL of cholesterol (5 mg/mL in ethanol) per 1 L). Static-free polyethylene microsphere beads (WPMS-1.00 ɸ250-300 μm, Cospheric LLC) were added at 10%. Thirty fertilized eggs were extracted from young adults and introduced into the solution. The bags were sealed 1.5 d before the SpaceX Crew-8 launch and transported to ISS Kibo. After 8 d, some bags were used to observe F1 adult motility using a high-definition camera. The remaining bags were frozen below -80°C in orbit. Space flight samples averaged 21.3°C (min -1.6°C, max +3.6°C), whereas ground control samples averaged 22.1°C (min -2.3°C, max + 0.4°C).

Culture bags with syringe ports (Fig. 1B) contained 200 L2 larvae from four GFP-transgenic strains in 12 ml of solution. After 6.7 days of preparation and arrival at the ISS at 11.7 ± 0.6°C, bags were incubated under 1G centrifugation in CBEF and at μG, at 21.4°C (min -0.4°C, max +0.1°C). Three days later, F1 progenies reached the L4 stage; 6 ml was transferred to a new bag with 6 ml M9C E. coli medium (OD600=4.) containing FUdR (75 μM) to prevent mixing. The culture was continued for 9.5 d until the D10 stage. Then, 6 ml was transferred to an empty bag and frozen below -80°C in MELFI freezer within ISS Kibou. For the remaining 6 ml on day 4.7 (D01, L4 larvae) and day 14 (D10, aged adults), 15 ml of M9 buffer with 0.99% PFA was added, fixed, and stored at 4°C. The samples were returned to Tohoku University, where frozen samples were used for gene expression analysis and PFA-fixed samples were used for GFP observation.

### Expression analyses

Approximately 1,000 worms (L4 to young adult cohort) were collected by thawing frozen samples on ice and washing them with ice-cold M9 buffer to remove eggs, small larvae, and debris. RNA extraction and sequencing were performed using the MGIEasy RNA Library Prep Set, MBIEasy Circularization Kit, DNBSEQ-G400RS Sequencing Kit, and DNBSEQ-G400 (MGI Tech).

### Body length measurement of space-flown adults

Twenty random adults were selected from approximately 500 frozen orbital nematodes using platinum wire under a stereomicroscope. Body length was measured using a DP74 microscopic digital camera system (Olympus, Tokyo, Japan) 7-9.

### Imaging and quantitative analysis

D01 worm movements were recorded in orbit using a high-resolution camera, and swimming behavior frequency was calculated from videos transmitted from the JAXA Kibo ISS module. Movement was also captured using confocal space microscopy (COSMIC), and images were downlinked. Representative images are shown in Supplementary Movie 1.

Space samples were removed from plastic bags using syringes and washed with M9 buffer prior to imaging. Transgenic strains with fluorescent dopaminergic neurons, GABAergic D-type motor neurons, mitochondria, and mitochondrial Ca^2+^ signals were observed using a scanning confocal microscope (Olympus FV10i-ASW). Animals were stained with rhodamine-conjugated phalloidin to observe the muscle fibers 31. For CEP neurons, blebs in the dendrites were measured 33. For D-type motor neurons, blebs and axonal defects in the mid-body area, including the commissural axons of VD7, DD4, VD8, and VD9, were counted (Supp Fig 1) 52. Blebs in four axon commissures were categorized as 1–5, 6–10, 11–15, and ≥ 16, and the proportions were plotted on a stacked graph.

Synaptic structures were examined using a Zeiss LSM 700 confocal microscope. SNB-1::GFP presynaptic puncta were analyzed in the dorsal nerve cord anterior to vulva (*unc-25p::SNB-1::GFP*) and nerve ring (*mec-7p::SNB-1::GFP*). For ALM neurons, images were captured at a 12-bit resolution with a maximum gray level of 4095. GLR-1::GFP postsynaptic puncta were analyzed along the ventral nerve cord, anterior to the vulva. The number and size of fluorescent puncta were analyzed using ImageJ software.

### Neuromuscular aging observation (ground experiment)

To inhibit egg laying during aging experiments, NGM plates with FUdR (50 μM) were covered with M9C liquid medium containing *E. coli* OP-50, and 30-50 L4 nematodes were placed on the plates. Adult worms were moved to new FUdR+M9 plates every three days, with dead nematodes removed daily. To observe GABAergic D-type motor neurons, adult worms on days 1, 10, and 15 were washed with M9 buffer and observed under an Olympus BX50 microscope. Mitochondria in the body wall muscles of 3-, 7-, and 10-day-old adults were observed using scanning confocal microscopy. Samples were mounted on a 1% agarose pad and paralyzed with levamisole (motor neurons) or NaN_3_ (mitochondrial observation).

### Imaging of live nematode moving on the ground and simultaneous imaging of muscle cytoplasmic calcium

Using ATU2301 and ATU2311 *mec-4* strains with the *goeIs3* transgene, cytoplasmic Ca^2+^ levels of adult hermaphrodites during crawling on *E. coli* OP50 plates were captured using a stereomicroscope and digital camera. Videos were converted to Royal Color and time-lapse images using the ImageJ software.

### Statistical Analysis

Statistical analysis was performed using an unpaired two-tailed Student’s t-test or one-way analysis of variance (ANOVA) with Tukey’s post-hoc analysis. Microsoft Excel, Prism software and Chi-square test using BellCurve were used for statistical testing. The tests are indicated in the figure legends. Statistical significance was set at p<0.05. Similar letters between groups indicate no significance, whereas different letters indicate significant differences.

## Data availability

All RNA-seq global gene expression data were deposited in the DDBJ BioProject (PRJDB37534). Any additional information required to reanalyze the data reported in this study is available from the lead contact upon request.

## Acknowledgments

We thank Koichi Wakata for conducting the NIS spaceflight experiment on the ISS. NIS was supported by JAXA, and MME 2 was supported by the UK and ESA space agencies. This experiment was supported by JAXA’s Cell Biology Experiment Project and JSPS KAKENHI grants 25H01374 and AMED-CREST (16814305) (AH) and AMED-Moonshot (JP22zf0127001, 24zf0127001h0004); NRF grants 2021R1A2C101178312 and RS-2024-00409403; Asian Office of Aerospace Research and Development FA2386-24-1-4002; BBSRC grants BB/N015894/1 and BB/P025781/1; UK Space Agency grant ST/R005737/1; and NIH NIAMS ARO54342.

C. elegans strains were provided by the Caenorhabditis Genetics Center (NIH P40 OD010440).

## Author contributions

A.H. designed the study and supervised the spaceflight experiment. J.I.L., T.E., N.J.S., and Ak.H. were co-investigators. JH.M., JI.H., N.H., T.H., Y.H., I.S., J.I.L., and A.H. conducted various experiments. A.A.A., K.O., M.U., A.V.AJr., Bs.K., Ak.H. T.A. supported some experiments. A.H. and J.I.L. wrote the manuscript. All the authors approved the final manuscript.

## Code availability

Not applicable

---

**Supplementary Table 1. GO enrichment analysis of genes with decreased expression in the D01 wild type grown under μG conditions compared to the terrestrial 1G environment**

**Supplementary Table 2. GO enrichment analysis of genes with increased expression in the D01 wild type grown under μG conditions compared to the terrestrial 1G environment**

**Supplementary Table 3. GO enrichment analysis of genes with increased expression in the D01 wild type grown under μGB conditions compared to the μG environment**

**Supplementary Table 4. GO enrichment analysis of genes with decreased expression in the D010 wild type grown under 1G conditions compared to the D01 worms.**

**Supplementary Table 5. GO enrichment analysis of genes with increased expression in the D10 wild type grown under 1G conditions compared to the D01 worms.**

**Supplementary Table 6. GO enrichment analysis of genes with decreased expression in the D10 wild type grown under μG conditions compared to 1G conditions.**

**Supplementary Table 7. GO enrichment analysis of genes with increased expression in the D10 wild type grown under μG conditions compared to 1G conditions.**

**Supplementary Table 8. GO enrichment analysis of genes with decreased expression in the D10 wild type grown under μGB conditions compared to 1G conditions.**

**Supplementary Table 9. Fold changes in gene expression of D01 wild type and *mec-4* mutants compared between 1G vs. μG and μG vs. μGB.**

**Supplementary Fig. 1. Age-related axonal neurodegeneration in D-type GABAergic neurons on the ground.** (A) Diagram showing D-type GABAergic motor neurons (green) in the *C. elegans* strain CZ13799 (*juIs76* [*unc-25p::GFP* + *lin-15*(*+*)] II) with GFP in GABAergic neurons. Dorsal side is top. The analyzed axonal commissures are indicated in red. (B) Fluorescence images showing axonal defects in GABAergic motor neuron commissures: “Join and reach” (top left), “Turn and extend” (top right), “Stop” (bottom left), and “Branched” (bottom right), marked with white arrows. Scale bar: 100 µm. (C) Analysis of axonal defects in the commissures is shown as box-and-whisker plots (median, interquartile range, 10-90 percentile). Different letters indicate significant differences (*p < 0.05). Sample sizes: n =2 5, 42, and 35. (D) Images showing bleb formation in the commissural axons, marked by yellow arrowheads. Scale bar: 100 µm. (E) Damage assessment of GABAergic neurons in space-grown C. elegans: blebs in four axon commissures categorized as 1–5, 6–10, 11–15, and ≥ 16, plotted as the proportion of individuals. The same individuals as in (C) were used. 3 × 4 chi-square test for significance. *p-value: <0.05.

**Supplementary Fig. 2. Microgravity-induced bleb formation in dopaminergic CEP neurons of *C. elegans*.** (A) Fluorescence images of transgenic *C. elegans* TG2435 (*vtIs* [*dat-1p::GFP* + *rol-6*(*su1006*)] V) expressing GFP in dopaminergic CEP neurons. Arrowheads show bleb formation in CEP neurons, and arrows indicate neuronal cell bodies. Scale bar: 50 μm. (B) To assess GABAergic motor neuron damage in space-grown C. elegans, blebs in four axon commissures were categorized as 1–5, 6–10, 11–15, and ≥ 16, with proportions plotted on a stacked graph. Sample sizes from left to right: n = 19, 20, 27, 18, 17, and 25. A 6×4 chi-square test was performed. *p<0.05, **p<0.01.

**Supplementary Fig. 3. Muscle mitochondrial morphology in D01 L4 larvae grown under different gravity conditions.** Representative images of body wall muscle cells from two D01 L4 larvae, ATU3301 (*ccIs4251* [*myo-3p::GFP::LacZ::NLS* + *myo-3p::mitochondrial GFP*+*dpy-20* (*+*)] and *aceIs1* [*myo-3p::mitochondrial LAR-GECO* + *myo-2p::RFP*]), cultured under space μG, μGB, and artificial 1G. The scale bar represents 10 μm.

**Supplementary Fig. 4. Classification of age-related muscle mitochondrial damage.** Mitochondrial age-related changes were classified by their morphology according to Supplementary Figure 4 as follows: “intermediate,” showing a combination of an interconnected mitochondrial network and some small fragmented mitochondria; “fragmented,” showing mostly small fragmented mitochondria; “very fragmented,” showing small, round mitochondria; “swelling,” showing sparse, small, swollen mitochondria; and “severely swollen,” showing severely swollen mitochondria, with significant loss of GFP signal and progressive nuclear breakdown. The scale bar represents 10 μm.

**Supplementary Fig. 5. Volcano plot and GO analyses of gene expression changes in *mec-4* mutants under terrestrial 1G conditions.** (A) Volcano plot analysis comparing gene expression by RNA sequencing of the DO1 cohort (L4 larvae to young adults) in wild-type and mec-4 mutants at 1G. Expression ratio log2(FC) and -log10FDR were plotted using three biological replicates. Blue: decreased genes, orange: increased genes, gray: not significant, green: cuticle/collagen genes (blue/orange outlines: increased/decreased), yellow: autophagy genes (red outlines: increased). (E) GO enrichment analysis of 291 genes decreased in *mec-4* mutants vs. wild type. (F) GO enrichment analysis of 411 genes increased in *mec-4* mutants vs. wild type.

**Supplementary Movie 1. The movement behavior of D01 worms was recorded using confocal space microscopy (COSMIC) and a high-resolution camera at the JAXA Kibo experimental module on the ISS, as well as using both ground-based models.** The acquired images were converted to mp4 format using Image J and MS PowerPoint software.

**Supplementary Movie 2. Imaging of live moving wild type and *mec-4* (*u253*) mutant adults and simultaneous muscle cytoplasmic calcium imaging on the ground. The acquired videos were converted to Royal Color using ImageJ and MS PowerPoint software.**

